# Direct exposure to SARS-CoV-2 and cigarette smoke increases infection severity and alters the stem cell-derived airway repair response

**DOI:** 10.1101/2020.07.28.226092

**Authors:** Arunima Purkayastha, Chandani Sen, Gustavo Garcia, Justin Langerman, Preethi Vijayaraj, David W. Shia, Luisa K. Meneses, Tammy M. Rickabaugh, A. Mulay, B. Konda, Myung S. Sim, Barry R. Stripp, Kathrin Plath, Vaithilingaraja Arumugaswami, Brigitte N. Gomperts

## Abstract

Most demographic studies are now associating current smoking status with increased risk of severe COVID-19 and mortality from the disease but there remain many questions about how direct cigarette smoke exposure affects SARS-CoV-2 airway cell infection. We directly exposed mucociliary air-liquid interface (ALI) cultures derived from primary human nonsmoker airway basal stem cells (ABSCs) to short term cigarette smoke and infected them with live SARS-CoV-2. We found an increase in the number of infected airway cells after cigarette smoke exposure as well as an increased number of apoptotic cells. Cigarette smoke exposure alone caused airway injury that resulted in an increased number of ABSCs, which proliferate to repair the airway. But we found that acute SARS-CoV-2 infection or the combination of exposure to cigarette smoke and SARS-CoV-2 did not induce ABSC proliferation. We set out to examine the underlying mechanism governing the increased susceptibility of cigarette smoke exposed ALI to SARS-CoV-2 infection. Single cell profiling of the cultures showed that infected airway cells displayed a global reduction in gene expression across all airway cell types. Interestingly, interferon response genes were induced in SARS-CoV-2 infected airway epithelial cells in the ALI cultures but smoking exposure together with SARS-CoV-2 infection reduced the interferon response. Treatment of cigarette smoke-exposed ALI cultures with Interferon β-1 abrogated the viral infection, suggesting that the lack of interferon response in the cigarette smoke-exposed ALI cultures allows for more severe viral infection and cell death. In summary, our data show that acute smoke exposure allows for more severe proximal airway epithelial disease from SARS-CoV-2 by reducing the mucosal innate immune response and ABSC proliferation and has implications for disease spread and severity in people exposed to cigarette smoke.

## INTRODUCTION

Coronavirus disease (COVID-19) is an infectious disease caused by the newly discovered severe acute respiratory syndrome-related coronavirus 2 (SARS-CoV-2) that is responsible for the current pandemic that is endangering lives globally. SARS-CoV-2 is an enveloped positive-sense single-stranded RNA virus that enters its host cell by binding to the angiotensin converting enzyme 2 (ACE2) receptor (Ziegler et al., 2020). SARS-CoV-2 primarily targets the respiratory tract and ACE2 is expressed in a gradient along the airways with higher expression from proximal to distal and is present on ciliated cells and some secretory cell subtypes (Hou et al., 2020). A low percentage of COVID-19 cases develop pneumonia with diffuse alveolar damage which can develop into acute respiratory distress syndrome (ARDS) (Lai et al., 2020). Severe COVID-19 lung disease has been most closely associated with older age, especially age over 65 years. Among these older hospitalized adults, underlying medical conditions have been associated with severe COVID-19, which include hypertension, obesity, diabetes mellitus and chronic lung disease (Sanyalou et al., 2020).

The Surgeon General’s Report in 1964 determined that cigarette smoking is the most important cause of chronic bronchitis and emphysema, in addition to causing lung cancer. Since then, smoking has been found to cause Idiopathic Pulmonary Fibrosis (IPF) and asthma as well as being the primary cause of chronic obstructive pulmonary disease (COPD)(National Center for Chronic Disease Prevention and Health Promotion (US) Office on Smoking and Health, 2014). Therefore, cigarette smoking is one of the most important and common causes of chronic lung disease. Mechanistically, smoking has been shown to reduce mucosal innate immunity leading to increased viral replication. One of the underlying mechanisms is degradation of the Type I interferon receptor (HuangFu et al., 2008) and another mechanism involves reduction of the immune response in the airways by inhibiting type II interferon-dependent gene expression through a decrease in Stat1 phosphorylation ((El-Mahdy et al., 2009). Another study found that cigarette smoke extract reduces the immediate-early, inductive, and amplification phases of the type I IFN response that was abrogated with glutathione antioxidant treatment (Bauer et al., 2008).

Given the importance of cigarette smoking in the development of chronic lung diseases, it has been suggested that smoking may be a significant risk factor for severe COVID-19. Accordingly, the World Health Organization reviewed the available evidence and concluded that smoking is associated with increased severity of disease and death in hospitalized COVID-19 patients, although they could not quantify the risk to smokers (Igić, 2020). The lack of clarity on the issue is likely because there has been a lower than expected prevalence of smoking reported in retrospective and observational databases because of incomplete reporting of smoking status in patients in emergency situations. Therefore, some studies reported no increase in smoking-related disease whereas more in depth analyses of demographic data showed an increased risk of severe COVID-19 associated with smoking (Guo, 2020)(Vardavas and Nikitara, 2020)(Zhao et al., 2020). A recent study showed that smoking is a risk factor for more severe COVID-19 among young adults (Adams et al., 2020).

Several studies have examined ACE2 expression in the airways of smokers. Zhang et al looked at bulk RNA-sequencing and single cell RNA sequencing (RNA-seq) from the proximal and distal airways and found that smoking increased ACE2 expression in the distal airway (Zhang et al., 2020). Similarly, another group also examined multiple airway datasets and found upregulation of pulmonary ACE2 gene expression in all patients who had ever smoked compared with nonsmokers, irrespective of tissue type or COPD status (Cai et al., 2020). However, there have been no direct studies to examine the effect of smoking on the airway epithelium in the setting of SARS-CoV-2 infection and therefore it has remained unclear as to whether smoking influences SARS-CoV-2 infection.

Therefore, we developed a system to expose primary human mucociliary epithelial cultures at the air-liquid interface to cigarette smoke and subsequently infected the culture with live SARS-CoV-2. This system allowed us to uncover the complex cellular and molecular interactions of the co-morbidities of smoking and viral infection. We found an increase in the number of infected cells after cigarette smoke exposure and that acute cigarette smoke exposure increases airway basal stem cells (ABSCs) while SARS-CoV-2 prevents the normal repair response from ABSCs. We also found that SARS-CoV-2 infection downregulates many host genes in all airway cell types and upregulates the interferon response, which could account for the relatively low number of infected cells in the ALI cultures. Short term cigarette smoke exposure reduces the interferon response, suggesting that the modulation of the interferon response by SARS-COV-2 is causally linked to higher infection in smoke-exposed cultures. Consistent with this hypothesis we found that the smoke-induced increase in SARS-COV-2 infection could be abrogated by treatment with exogenous interferonβ-1.

## RESULTS

### Cigarette smoke exposure increases SARS-CoV-2 infection in the airway epithelium and SARS-CoV-2 prevents the stem cell-mediated repair response

ALI cultures are derived from primary human ABSCs and when differentiated on a transwell membrane in specific media demonstrate the mucus, ciliated and secretory cells of the proximal airway epithelium (Supplemental Table S1)(Gruenert et al., 1996). As the surface of the cultures is directly in contact with air, we are able to directly expose the cells to cigarette smoke. We found that exposure of ALI cultures to cigarette smoke for three minutes once a day for four days was sufficient to induce airway injury and repair with an increased number of ABSCs but resulted in minimal cell death (data not shown). Consequently, we used ABSCs from a healthy lung transplant donor for ALI cultures and exposed them to short cigarette smoke intervals for 4 days before infecting with SARS-CoV-2 (MOI = 0.1) as shown in the schematic in Figure 1A. Control ALI cultures received no infection (mock infection) or smoke exposure, infection alone or smoke exposure alone. Three days after infection or mock treatment, we examined the cultures for evidence of SARS-CoV-2 replication. We performed quantitative real time PCR (q-RT-PCR) on RNA obtained from the airway epithelial cells and found an increase in viral load after cigarette smoke exposure compared to infection alone (Figure 1B, Supplemental Table S2). Employing immunofluorescent staining (IF) to detect SARS-CoV-2 dsRNA, we found that there was an increased number of infected cells in ALI cultures that had smoke exposure (p<0.001) (Figure 1C,D). We quantified the number of ciliated and mucous airway epithelial cell types across all conditions by immunostaining with cell-type specific markers (Figure 1C-F). We found that smoke exposure alone increased the number of Muc5AC-expressing mucus cells and that there was a decrease, albeit not significant, in the number of ciliated cells marked by the presence of acetylated β-tubulin (Figure 1C-F). We found that the number of cells expressing acetylated β-tubulin or Muc5AC did not significantly change with SARS-CoV-2 infection (Figure 1C-F). We then assessed the number of ABSCs under these conditions by immunostaining for keratin 5 (KRT5), which is the key stem cell type orchestrating the repair response by proliferation. We found that smoking increased the number of ABSCs as part of a repair response but that SARS-CoV-2 viral infection alone or viral infection with smoking did not trigger the expected increase in ABSCs needed for repair (Figure 1G,H). Furthermore, immunostaining for Ki67 combined with PCNA to identify all proliferating ABSCs showed that the ABSCs were induced to proliferate after smoke exposure but that SARS-CoV-2 infection alone or the combination of smoke exposure and infection did not significantly alter the proliferation rate of ABSCs, implying that SARS-CoV-2 inhibits the repair process in the airway epithelium (Figure 1I,J). Immunostaining for cleaved caspase 3 (CC3) revealed that apoptosis was infrequently seen in the ALI cultures at 3 days post infection or with cigarette smoke exposure alone, but the combination of cigarette smoke and infection significantly increased the number of apoptotic cells (Figure 1K,L). Higher magnification revealed that most but not all the apoptotic cells were infected with SARS-CoV-2 and that not all infected cells were undergoing apoptosis. SARS-CoV-2 infection is therefore promoting cell death while reducing the normal airway epithelial repair response. As ACE2 is the receptor for SARS-CoV-2, we examined ACE2 expression by immunostaining and found that there was a trend towards increased ACE2 expression after cigarette smoke exposure and that there was no change in ACE2 expression in infected cells or infected cells exposed to cigarette smoke (Figure 1M,N).

**Figure 1.**
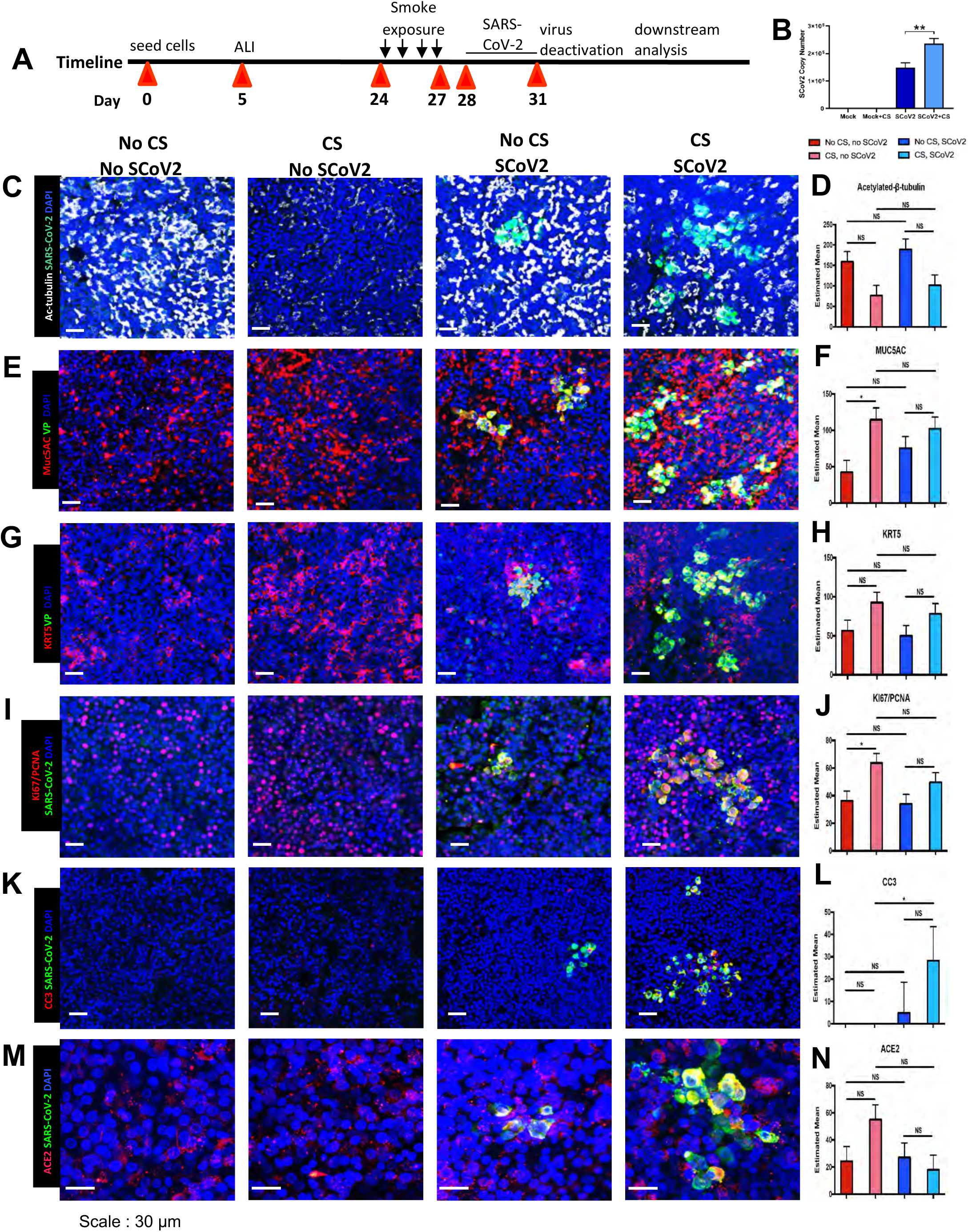
Cigarette smoke exposure increases SARS-CoV-2 infection in the airway epithelium. **A.** Experimental schematic outline showing the total days in culture with days of experimental manipulations **B.** Quantification of SARS-CoV2 viral load with and without smoking exposure by quantitative real-time PCR of viral mRNA **C.** IF images of ciliated cells (white) and SARS-CoV-2 (green) infected cells in ALI cultures across the four exposure conditions of no smoking and no virus, smoking and no virus, no smoking and virus and both smoking and virus exposures **D.** Quantification of number of ciliated cells across the four exposure groups **E.** IF images of Muc5AC (red) mucus cells and SARS-CoV-2 (green) infected cells in ALI cultures across the four exposure conditions **F.** Quantification of number of Muc5AC+ mucus cells across the four exposure groups **G.** IF images of K5 (red) ABSCs and SARS-CoV-2 (green) in ALI cultures across the four exposure conditions **H.** Quantification of number of ABSCs across the four exposure groups **I.** IF images of ACE2 (red) and SARS-CoV-2 (green) in ALI cultures across the four exposure conditions **J.** Quantification of number of ACE2+ cells across the four exposure groups **K.** IF images of both Ki67 and PCNA (red) and SARS-CoV-2 (green) expressing cells in ALI cultures across the four exposure conditions **L.** Quantification of number of proliferating cells across the four exposure groups **M.** IF images of cleaved caspase 3 (CC3)(white) and SARS-CoV-2 (green) for apoptosis across the four exposure conditions **N.** Quantification of number of apoptotic cells across the four exposure groups Bar graph represents SEM, n = 3-6. *p < 0.05, ** p<0.01, ns = not significant by Turkey test.

### Smoking reduces the innate immune response in SARS-CoV-2 infection and interferon treatment abrogates infection after cigarette smoke exposure

We then sought to determine a possible mechanism for the higher cellular infection seen upon exposure of the ALI culture with both smoking and SARS-CoV-2. To explore the differences in SARS-CoV-2 infectivity between cigarette smoke exposure and no exposure, we applied single cell RNA-seq to determine the transcriptional alterations taking place which might explain the differences in SARS-CoV-2 infectivity between cigarette smoke exposure and no exposure. We used the same experimental timeline as outlined in the schematic in Figure 1A and pretreated cells with/without cigarette smoke and then exposed the ALI cultures to SARS-CoV-2 or mock treatment for three days before single cell dissociation for RNA-seq. From these four conditions, we recovered 19361 single cell transcriptomes that passed filtration criteria. Upon plotting the data, we observed that mock-treated cells and cells exposed to cigarette smoke were largely mixed and that cells with SARS-CoV-2 infection were separated (Figure 2A), indicating that SARS-CoV-2 infection induces profound transcriptional changes. Despite these differences, we detected all major expected cell types of the lung epithelium within the ALI cultures (Figure 2B), expressing typical human airway cell type marker genes (DNAAF, ciliated cells; KRT5, basal cells; and MUC5B secretory cells; Figure 2C). We detected SARS-CoV-2 transcripts in all major cell types and found the proportion of infected cells to be highest in FOXN4+ cells (Supplementary Figure 1A).

**Figure 2.**
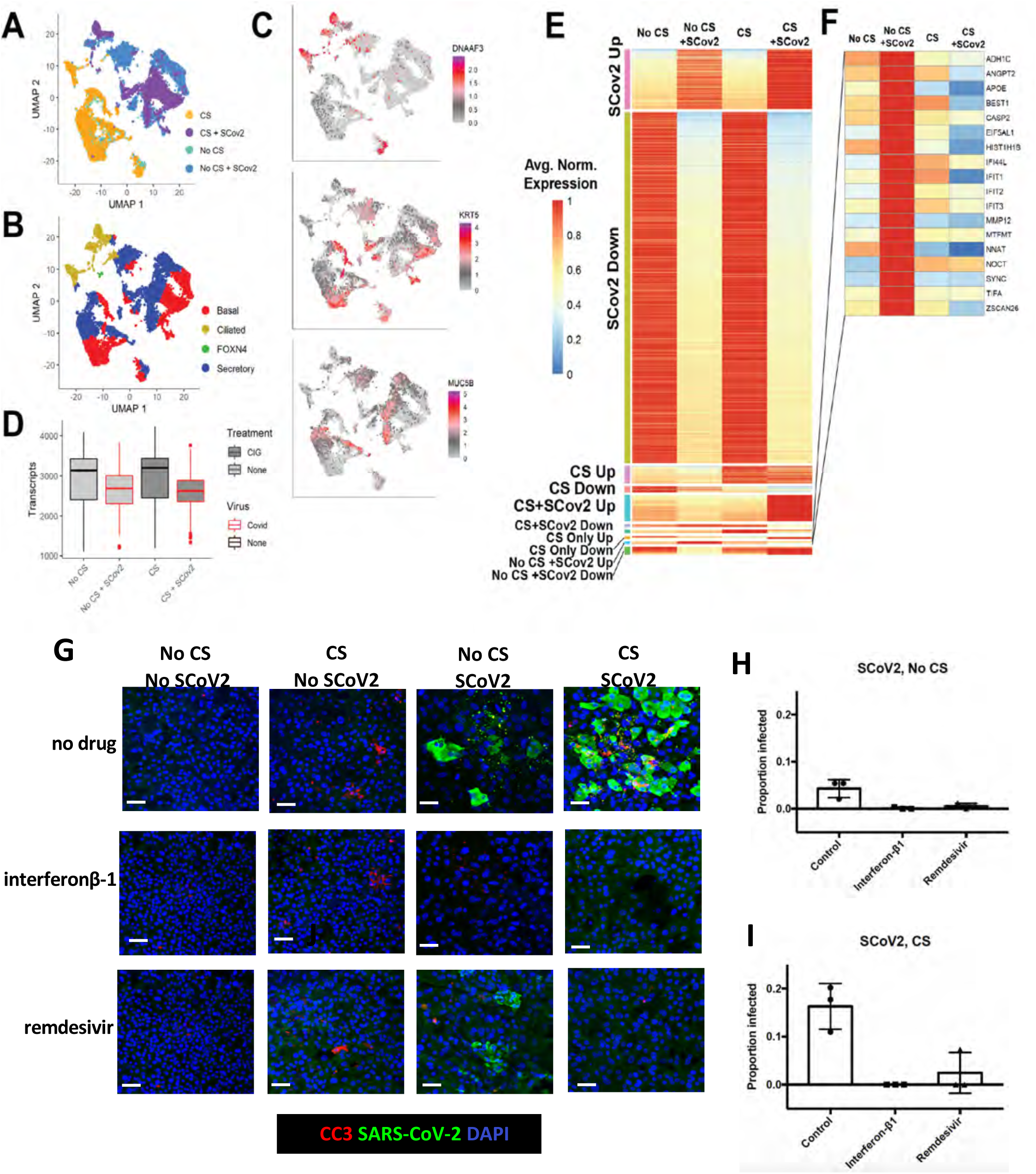
Smoking reduces the Interferon response in SARS-CoV-2 infection and Interferon abrogates infection after cigarette smoke exposure. **A.** Reduced dimensionality graph showing the position of 19361 cells exposed to no cigarette smoke or virus, cigarette smoke (CS) alone or virus (SARS-CoV-2) alone or cigarette smoke and virus. Cells are identified as shown in key **B.** On the same plot as in (A), cell types are shown based on the expression of known human airway genes. Cell type is colored as shown in key. **C.** Shown are cells colored by their normalized expression value of cell type marker genes *Dnaaf3* (ciliated cells), *Krt5* (basal cells), and *Muc5b* (secretory cells). **D.** Boxplots which capture the distribution of normalized transcripts in all cells per sample, as separated by cigarette and SARS-CoV-2 exposure. **E.** Heatmap showing the percent expression of all genes found to be differentially expressed in at least one condition. CS indicated cigarette smoke treatment and SCov2 indicates SARS-CoV-2 viral exposure. **F.** Popout of heatmap showing the specific genes which are induced by SARS-CoV-2 but not in cigarette smoke exposure with SARS-CoV-2 (No CS + SCov2 Up). Bar graph represents SEM, n = 3-6. ∗p < 0.05, ** p < 0.01, *** p < 0.001 by Student’s t test. **G.** IF images of SARS-CoV-2 (green) infected cells across the four exposure groups with no drug treatment, interferonβ-1 or remdesivir treatment. **H.** Quantification of the number of infected cells with each treatment in the four exposure groups **I.** Quantification of the number of apoptotic cells with each treatment in the four exposure groups Bar graph represents SEM, n = 3-6. *p < 0.05, ** p<0.01, *** p<0.001. ns = not significant by Student’s t test.

Interestingly, the largest difference among samples was the global downregulation of genes in SARS-CoV-2 exposed samples, with approximately 10-15% fewer transcripts in exposed cells compared to uninfected controls (Figure 2D). Using differential expression to determine genes specific to each condition, we found that the largest change in the data was in fact the downregulation of 2805 genes in SARS-CoV-2 exposed samples and this was consistent across all major cell types (Figure 2E, Supplementary Figure 1B). Among the downregulated genes were genes related to the viral immune response, apoptotic signaling, and various metabolic processes. 475 genes were upregulated upon exposure to the virus, including genes related to interferon signaling and chromatin organization. Although the majority of gene expression changes were controlled by virus exposure, we found another 559 genes that were altered in response to cigarette smoke exposure. Even prior to virus exposure, cigarette smoke downregulated the innate immune response and stimulated airway differentiation genes. In response to SARS-CoV-2, cigarette smoke-treated cells downregulated genes related to metabolic and wound healing processes and upregulated cilia-related genes. Interestingly, a small class of genes was induced in the non-cigarette-exposed cells in response to SARS-CoV-2 but downregulated in the cigarette-exposed infected ALI cultures. These genes included Interferon Induced Protein With Tetratricopeptide Repeats (IFIT) *IFIT1, IFIT2, IFIT3* and Interferon Induced Protein 44 Like (*IFI44L)* and indicated an interferon mediated response induced by viral infection, that was repressed below the level in control cells in infected cigarette-treated cells (Figure 2F, Supplemental Table S3). These data suggest that cigarette smoke exposure prevents an effective interferon-based response to the SARS-CoV-2 virus.

Because the interferon response was increased in viral infection alone but decreased in the setting of smoking with viral infection, we reasoned that the greater infection seen upon treatment with cigarette smoke and SARS-CoV-2 was, at least in part, due to smoking induced reduction in the innate immune response. We therefore treated the cigarette exposed or unexposed and virally infected or uninfected ALI cultures with interferonβ-1 and included remdesivir (a direct acting antiviral agent) or no drug, as controls. These were added to the ALI cultures after cigarette exposure and just prior to viral infection and were present in the cultures for three days after which the cultures were fixed and immunostained for the apoptotic marker CC3 and SARS-CoV antigen. We found that interferonβ-1 completely abrogated the infection (Figure 2G-I), while remdesivir also showed an inhibitory response to viral infection. This shows that a lack of interferon response is at least one important mechanism for the higher SARS-CoV-2 infectivity seen in smoking exposed ALI cultures.

## DISCUSSION

There has been some controversy about whether cigarette smoke exposure increases SARS-CoV-2 infectivity and whether this might lead to more severe disease. Our primary human mucociliary epithelial cell culture model with direct exposure to cigarette smoke and SARS-CoV-2 infection demonstrates that even short-term exposure to cigarette smoke in primary human airway cells from a previously healthy patient who was not a chronic smoker leads to increased infection. Our data suggests three potential and synergistic mechanisms for this: reduced innate immunity of the cells leading to greater infection, lack of ABSCs to appropriately proliferate for airway repair after infection, which could lead to worse tissue damage and/or stem cell exhaustion and increased apoptosis in airway cells exposed to cigarette smoke and SARS-CoV-2.

The reduction in the interferon response by smoking is one mechanism whereby SARS-CoV-2 may more easily enter and replicate in epithelial cells. Interestingly, interferon response genes were actually induced by the virus, which may explain why we only see a low percentage of infected cells in the ALI cultures and why the majority of the human population does not develop severe respiratory infections. Our data suggest that there are additional factors from cigarette smoke exposure that could make the cells more vulnerable to infection. For example, the synergistic increase in apoptotic cells after cigarette smoke exposure and infection suggests that the cigarette smoke may act with the virus to potentiate other pathways, such as the DNA damage response pathway, to injure the epithelium. We also saw a reduction in cell division in the ABSCs which prevented the normal host response with repair and regeneration of the airway. This result was in contrast to smoking injury where ABSCs proliferate robustly to repair the airway epithelium. Despite smoking injury, the SARS-CoV-2 response was dominant and prevented the normal ABSC proliferation response.

The interferon response is reported to be reduced or delayed in serum levels in ferrets and patients with severe COVID-19 where pro-inflammatory cytokines and chemokines are more dominant (Blanco-Melo et al., 2020). Others have found a temporal expression of interferon genes after SARS-CoV-2 infection (Yoshikawa et al., 2010). We noted an increase in interferon response genes in ALI cultures at 3 days post infection with low MOI (0.1). This may reflect the fact that the model does not include inflammatory cells but does suggest that an intact, normal airway epithelium could function as a barrier to COVID-19 with a good innate immune response and could explain why the majority of infected individuals have mild symptoms and even asymptomatic carriage. Abrogation of SARS-CoV-2 infection with type I interferons has been reported by others in other settings (Lokugamage et al., 2020; Mantlo et al., 2020), including a small clinical trial (Wang et al., 2020).

The human primary ALI cell culture model provides a useful system for studying the direct effects of cigarette smoke on the airway epithelium. Our experiments directly examined the effects of short-term cigarette smoke exposure on the airway, which implies that current smokers are at risk of more severe infection. It is not clear whether former smokers will have the same risk of infection that current smokers do and this is something that remains to be tested. The increased number of infected cells in smokers has implications for more severe infection in smokers resulting in increased lung disease although our cultures are of the proximal airways so we could not directly assess for effects on diffuse alveolar damage or ARDS. Overall our data provides evidence for the need for aggressive health measures to stop smoking to reduce severe COVID-19.

## ACKNOWLEDGMENTS

We would like to thank Abdo Durra and Andrew Lund for their input on the manuscript. This work was supported by the NIH/NCI Grant R01CA208303 (BG), the Tobacco Related Disease Research Program (TRDRP) High Impact Pilot Research Award (HIPRA) 26IP-0036 (BG), the TRDRP HIPRA 29IP-0597 (BG), the UCOP Emergency Funding COVID19 TRDRP Seed grant Award R00RG2383 (BG), the Ablon Research Scholars Award (BG), the UCLA Oversight COVID-19 Research Committee (OCRC)(BG and VA), the UCLA Eli & Edythe Broad Center of Regenerative (BSCRC) (BG), the UCLA Medical Scientist Training Program grant (NIH NIGMS GM008042) (DS). KP was supported by the UCLA BSCRC, the David Geffen School of Medicine, NIH P01 GM099134, and a Faculty Scholar grant from the Howard Hughes Medical Institute. We appreciate the UCLA BSCRC Microscopy Core and the UCLA Translational Pathology Core Laboratory. This research was supported by NIH National Center for Advancing Translational Science (NCATS) UCLA CTSI Grant Number UL1TR001881. The following reagent was obtained through BEI Resources, NIAID, NIH: Polyclonal Anti-SARS Coronavirus (antiserum, Guinea Pig), NR-10361.

## AUTHOR CONTRIBUTIONS

**A**.**P**., **C**.**S**., **G**.**G**., **JL:** conception and design, collection and assembly of data, data analysis and interpretation,

**P**.**V**., **L**.**K**.**M**., **D**.**W**.**S**., **T**.**M**.**R**.: data analysis and interpretation and technical support.

**V**.**A**., **K**.**P**.: conception and design, data analysis and interpretation, manuscript writing,

**M**.**S**.**S**.: data analysis and interpretation

**B**.**R**.**S**., **A**.**M**., **B**.**K**.: conception and design

**B**.**N**.**G**.: conception and design, data analysis and interpretation, manuscript writing, final approval of the manuscript and financial support.

## STAR METHODS

### RESOURCE AVAILABILITY

#### Lead contact

Further information and requests for resources and reagents should be directed to and will be fulfilled by the Lead Contact, Brigitte N. Gomperts.

#### Materials Availability

This study did not generate new unique reagents.

#### Data and Code Availability

This study generated a single cell RNA sequencing dataset of ALI cultures that were exposed to cigarette smoke and/or SARS-CoV-2 or neither. These data are currently being uploaded into GEO. Accession number pending.

### Method Details

#### Human Tissue Procurement

Large airways and bronchial tissues were acquired from de-identified normal human donors after lung transplantations at the Ronald Reagan UCLA Medical Center. Tissues were procured under Institutional Review Board-approved protocols at the David Geffen School of Medicine at UCLA. For some experiments normal human bronchial epithelial cells (NHBE) were obtained from Lonza and all samples were de-identified.

#### ABSC Isolation

Human ABSCs were isolated following a previously published method by our laboratory (Hegab et al., 2012b, 2012a, 2014; Paul et al., 2014). Briefly, airways were dissected, cleaned, and incubated in 16U/mL dispase for 30 minutes at room temperature. Tissues were then incubated in 0.5mg/mL DNase for another 30 minutes at room temperature. Epithelium was stripped and incubated in 0.1% Trypsin-EDTA for 30 minutes shaking at 37°C to generate a single cell suspension. Isolated cells were passed through a 40μm strainer and plated for Air-Liquid Interface cultures.

#### Air-Liquid Interface (ALI) Cultures and In Vitro Treatments

24-well 6.5mm transwells with 0.4μm pore polyester membrane inserts were coated with 0.2mg/mL collagen type I dissolved in 60% ethanol and allowed to air dry. ABSCs were seeded at 100,000 cells per well in a 1:1 mixture of media:growth factor-reduced Matrigel to grow as tracheospheres or seeded directly onto collagen-coated transwells and allowed to grow in the submerged phase of culture for 4-5 days with 500μl media in the basal chamber and 200μl media in the apical chamber. ALI cultures were then established and cultured with only 500μl media in the basal chamber, and cultures were harvested at varying timepoints for IF studies. Media was changed every other day and cultures were maintained at 37°C and 5% CO_2_.

#### Tracheal Epithelial Cell (TEC) Plus and Serum-Free Media

Human ABSCs/NHBEs were grown in TEC Plus media and TEC serum-free media during the submerged and ALI phases of culture, respectively. TEC base media is DMEM/Ham’s F12 50/50 (Corning 15090CV). Table S1 indicates the media components and concentrations for TEC Plus and TEC serum-free media.

#### Cigarette Smoke *In Vitro* Treatments

A sterile chamber of 0.4 cu. ft. volume, with an attached vacuum pump, was used to generate and deliver cigarette smoke. The ALI plates (without lids) were placed inside the chamber and exposed to the smoke of a 1R3F research cigarette (University of Kentucky, Lexington, KY), which was attached to the vacuum chamber tube with a cigarette holder. The cigarette was burned 10% of its length and the smoke was introduced into the chamber via suction pump. A previously optimized treatment of 3-minute exposure/day was continued for 4 days.

#### SARS-CoV-2 infection

SARS-CoV-2, Isolate USA-WA1/2020, was obtained from Biodefense and Emerging Infectious (BEI) Resources of National Institute of Allergy and Infectious Diseases (NIAID). All the studies involving live virus were conducted in the UCLA BSL3 high-containment facility with appropriate institutional biosafety approvals. SARS-CoV-2 was passaged once in Vero-E6 cells and viral stocks were aliquoted and stored at -80°C. Virus titer was measured in Vero-E6 cells by TCID_50_ assay.

ALI cultures on the apical chamber of transwell inserts were infected with SARS-CoV-2 viral inoculum (MOI of 0.1; 100 µl/well) prepared in ALI TEC media. The basal chamber of the transwell contained 500 µl of ALI media. For mock infection, ALI media (100 µl/well) alone was added. The inoculated plates were incubated for 2 hr at 37 °C with 5% CO_2_. At the end of incubation, the inoculum was removed from the apical chamber. At selected timepoints live cell images were obtained by bright field microscopy. Striking cytopathic effect (CPE) was observed in SARS-CoV-2 infected cells, indicating viral replication and associated cell injury. At 72 hours post infection (hpi), viral infection was examined by immunocytochemistry (ICC) analysis using a SARS-CoV antibody [BEI Resources: NR-10361 Polyclonal Anti-SARS Coronavirus (antiserum, Guinea Pig). SARS-CoV antigen was detected in the cytoplasm of the infected cells, revealing active viral infection (Figure 1C,E,G,I,K,M).

#### Interferonβ-1 Drug Study

Once the ALI-SARS-CoV-2 infection system was established, we evaluated the effect of the interferonβ-1 (200ng/ml). remdesivir (10 µM), a well characterized, direct acting antiviral agent, was used as a control. The ALI cultures in 24-well plates were pretreated with Interferonβ-1 for 1 hour, then SARS-CoV-2 inoculum (MOI 0.1) was added. DMSO vehicle treated cells, with or without viral infections, and with and without smoking exposure were included as controls. At 72 hpi, the cells were fixed and immunostained with Polyclonal Anti-SARS-CoV to assess viral genome replication (Figure 1C).

#### Immunocytochemistry

ALI cultures were fixed in 4% paraformaldehyde for 15 minutes followed by permeabilization with 0.5% Triton-X for 10 minutes. Cells were then blocked using serum-free protein block (Dako X090930) for one hour at room temperature and overnight for primary antibody incubation. After several washes of Tris-Buffered Saline and Tween-20 (TBST), secondary antibodies were incubated on samples for 1 hour in darkness, washed, and mounted using Vectashield hardest mounting medium with DAPI (Vector Labs H-1500). IF images were obtained using an LSM700 or LSM880 Zeiss confocal microscope and composite images generated using ImageJ. The list of antibodies used is provided in the Key Resources Table as part of STAR Methods.

#### Quantitative Polymerase Chain Reaction (q-RT-PCR)

RNA was isolated with the RNeasy Mini Kit (Qiagen 74104) following manufacturer’s protocol and quantified using a NanoDrop Spectrophotometer (ThermoFisher). cDNA synthesis was performed using the TaqMan Reverse Transcription Reagents (ThermoFisher) or iScript cDNA Synthesis Kit (BioRad)s as indicated by the respective company. qPCR was then performed using the primers in Table S2. Samples were run in triplicate and fold changes in expression were determined using the comparative ΔC_T_ method and GAPDH was used as an endogenous control.

#### Single cell library generation and sequencing

Single cells were obtained by incubating ALI cultures in 500ul Accumax for an hour and 15 minutes. Cells were then fixed in cold methanol per the Illumina protocol and frozen at -80°C. Cells were then rehydrated in ice cold PBS and RNAse inhibitor per the Illumina protocol. Cells were captured using a 10X Chromium device (10X Genomics) and libraries prepared according to the Single Cell 3’ v2 or v3 Reagent Kits User Guide (10X Genomics, https://www.10xgenomics.com/products/single-cell/). Cellular suspensions were loaded on a Chromium Controller instrument (10X Genomics) to generate single-cell Gel Bead-In-EMulsions (GEMs). Reverse transcription (RT) was performed in a Veriti 96-well thermal cycler (ThermoFisher). After RT, GEMs were harvested, and the cDNA underwent size selection with SPRIselect Reagent Kit (Beckman Coulter). Indexed sequencing libraries were constructed using the Chromium Single-Cell 3’ Library Kit (10X Genomics) for enzymatic fragmentation, end-repair, A-tailing, adapter ligation, ligation cleanup, sample index PCR, and PCR cleanup. Libraries QC was performed by the Agilent Technologies Bioanalyzer 2100 using the High Sensitivity DNA kit (Agilent Technologies, catalog# 5067-4626) and quantitated using the Universal Library Quantification Kit (Kapa Biosystems, catalog# KK4824. Sequencing libraries were loaded on a NovaSeq 3000 (Illumina).

#### Data analysis

Raw sequencing data were filtered by read quality, adapter- and polyA-trimmed, and reads were aligned to a hybrid human hg38-Sars Covid2 transcriptome using the Cell Range software (10X Genomics) and the STAR aligner. Expression counts for each gene were collapsed and normalized to unique molecular identifiers to construct a cell by gene matrix for each library, filtered to keep cells with over 2000 transcripts and genes expressed in at least 0.05% of cells.

Data analysis was performed in R. Expression matrices were normalized by the total number of transcripts per cell in log space by dividing raw counts by the total number of transcripts per cell, then multiplying by 10,000. Two-dimensional visualization was obtained with the UMAP package. To identify major cell types in our normal integrated datasets, previously published lung epithelial cell type specific gene lists (Plasschaert et al., 2018) were used to create cell type-specific gene signatures, with cells assigned by maximal identity score. To identify differentially expressed genes between samples, we selected genes from every given pair of condition comparisons which satisfied an average expression difference of 33% either up or down in log normalized counts, filtered by a Bonferroni corrected p<0.01. Gene ontology enrichments were determined using the Metascape tool (Zhou et al., 2019). For heatmap generation, the mean expression per condition was calculated and then per gene normalized to the maximal sample value.

### Statistical Analysis

Data are presented as the mean ± standard error of the mean (SEM) values. A general linear model was used to assess the associations of smoking and virus to each outcome. The interaction of smoking and virus was evaluated in the model and multiple comparisons were performed using Tukey’s adjustment method. Calculated p-values are indicated on individual graphs. ∗p < 0.05, ∗∗p < 0.01, ∗∗∗p < 0.001, n.s. = not significant.

**Supplemental Table 1. MTEC medias - Related to STAR Methods**.

**Supplemental Table 2. Viral load PCR primers. Related to STAR Methods**

**Supplemental Table 3. List of upregulated and downregulated genes across the four exposure groups.**

**Related to Figure 2**

