## Supplemental Figure 1 for "Direct exposure to SARS-CoV-2 and cigarette smoke increases infection severity and alters the stem cell-derived airway repair response"

**Supplementary Figure 1.**

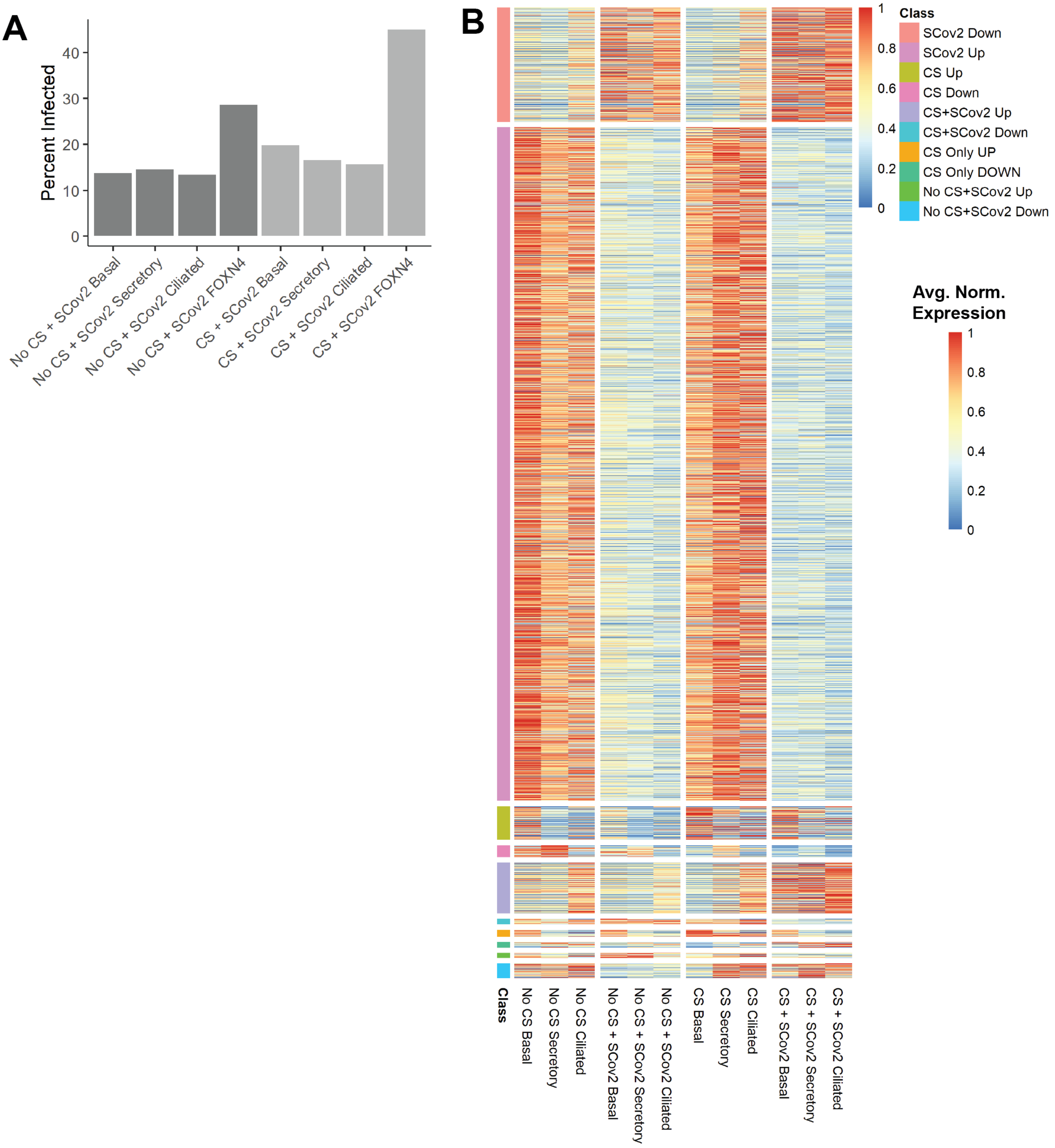

**Figure S1. Related to Figure 2.**

**A.** Bargraph showing the percent of cells in each condition per cell type which have detectable SARS-CoV-2 sequencing reads

**B.** Heatmap showing the differentially expressed genes from each condition as in Figure 2E, but now showing the average percent of gene expression across the three major cell types present in the culture. Expression normalized to maximal value per gene for condition and cell type combinations shown.
